# Application of Generalized Concentration Addition to Predict Mixture Effects of Glucocorticoid Receptor Ligands

**DOI:** 10.1101/2020.05.21.109439

**Authors:** Rosemarie de la Rosa, Jennifer J. Schlezinger, Martyn T. Smith, Thomas F. Webster

## Abstract

Environmental exposures often occur in complex mixtures and at low concentrations. Generalized concentration addition (GCA) is a method used to estimate the joint effect of receptor ligands that vary in efficacy. GCA models have been successfully applied to mixtures of aryl hydrocarbon receptor (AhR) and peroxisome proliferator-activated receptor gamma (PPARγ) ligands, each of which can be modeled as a receptor with a single binding site. Here, we evaluated whether GCA could be applied to homodimer nuclear receptors, which have two binding sites, to predict the combined effect of full glucocorticoid receptor (GR) agonists with partial agonists. We measured transcriptional activation of GR using a cell-based bioassay. Individual dose response curves for dexamethasone (full agonist), prednisolone (full agonist), and medroxyprogesterone 17-acetate (partial agonist) were generated and applied in three additivity models, GCA, effect summation (ES), and relative potency factor (RPF), to generate response surfaces. GCA and RPF yielded adequate predictions of the experimental data for two full agonists. However, GCA fit experimental data significantly better than ES and RPF for all other binary mixtures. This work extends the application of GCA to homodimer nuclear receptors and improves prediction accuracy of mixture effects of GR agonists.

## Introduction

Humans are exposed to multiple environmental contaminants and nonchemical stressors on a daily basis. The complexity of human exposures poses a challenge to risk assessment, which has traditionally evaluated individual chemicals (Carlin et al. 2013). Evaluating single chemicals is problematic and underestimates health risk since it does not account for potential mixture effects (Kortenkamp and Faust 2018). However, epidemiological studies are limited in their ability to assess mixture effects, and it is impractical to test all chemical combinations experimentally (Braun et al. 2016; Webster 2018). Consequently, alternative approaches are needed to address the mixture problem.

One approach is to predict mixture effects from individual dose-response curves with additivity models. This method requires defining a null hypothesis based on an assumed model (Rider et al. 2018). Independent action is a model traditionally applied to compounds with differing mechanisms of action. Alternatively, effect summation (ES) is often used for compounds with the same biological target and assumes that the joint effect is equivalent to the sum of the individual responses. ES is generally regarded as an inadequate model for evaluating mixtures because it allows predictions to exceed response boundaries and is only applicable to chemicals with linear dose-response curves (Berenbaum 1989). Concentration addition (CA) is another model used for compounds that act via similar mechanism, where the joint effect is estimated by the sum of each component scaled by their relative potency, which may in general depend on effect level. Silva et al. demonstrated the ability of CA to predict the additive effect of compounds with low potencies. The eight weakly estrogenic compounds tested in their study differed in relative potency but had similar dose-response shapes and efficacies, resulting in a special case of CA known as relative potency factor (RPF). However, CA and RPF cannot be applied to mixtures containing partial agonists since it assumes that all compound have the same maximum effect level.

Generalized concentration addition (GCA) addresses this limitation and allows mixture components to differ in efficacy (Howard and Webster 2009). Previous work demonstrates that GCA can be applied to mixtures of aryl hydrocarbon receptor (AhR) and peroxisome proliferator-activated receptor gamma (PPARγ) ligands (Howard et al. 2010; Watt et al. 2016). One requirement of GCA is specification of the dose-response function for each component in the model. For receptors with a single binding site, such as AhR and PPARγ, a Hill function with a coefficient of 1 is used to define the dose-response function. However, a different approach is required for receptors with two ligand-binding sites (e.g. homodimers), since the Hill coefficient is expected to exceed 1 and violate the invertibility requirement of GCA. For this reason, we used a pharmacodynamic concentration-response function that can be applied to receptors that homodimerize.

Steroid nuclear receptors are an important class of homodimer receptors that mediate the adverse effects of endocrine disrupting chemicals (Maqbool et al. 2016). Steroid receptors translocate from the cytoplasm to the nucleus after ligand binding and form homodimers that activate transcription. The glucocorticoid receptor (GR) is a steroid nuclear receptor expressed in nearly all human tissues and regulates transcription of 10-20% of genes in the human genome (Oakley and Cidlowski 2013). Glucocorticoid steroid hormones are endogenous GR ligands secreted in a circadian pattern and in response to stress (Biddie et al. 2012). Synthetic glucocorticoids also have been developed as anti-inflammatory and immunosuppressive drugs. The prevalence of long-term synthetic glucocorticoid usage in the United States is approximately 1.2% of the population (Overman et al. 2013). Environmental compounds, such as heavy metals and pesticides, are also capable of binding and modifying GR signaling (Odermatt and Gumy 2008; Gulliver 2017). Given the importance of this biological pathway and likelihood of concurrent exposure to GR ligands from multiple sources, the mixture effects of GR ligands warrant further investigation.

Here, we applied GCA to mixtures of GR ligands using a dose-response function for receptors that homodimerize. We used a cell line stably transfected with a glucocorticoid response element-dependent luciferase reporter to obtain individual dose-response curves for GR ligands, including two full agonists and a partial agonist. We also generated experimental data for binary mixtures of GR ligands to evaluate the response surface predictions generated by the GCA, ES and RPF additivity models.

## Materials and Methods

### Chemicals

Dexamethasone (DEX, cat. #D4902), prednisolone (PRED, cat. #P6004), and medroxyprogesterone 17-acetate (MPA, cat. #M1629) were all purchased from Sigma-Aldrich (St. Louis, MO).

### Measurement of GR activity (231GRE)

The 231GRE cell-based bioassay that we recently developed was used to measure plasma glucocorticoid levels (manuscript submitted for publication). Briefly, the MDA-MB-231 cell line was stably transfected with a luciferase reporter gene plasmid driven by three copies of a simple glucocorticoid-response element. 231GRE cells were cultured in Dulbecco’s Modified Eagle Medium (DMEM; Gibco, Waltham, MA) supplemented with 10% fetal bovine serum (FBS; Atlanta Biologicals, Atlanta, GA) at 37°C in an incubator with 5% CO_2_. Cells were switched to phenol red-free DMEM (Hyclone, Logan, Utah) containing charcoal-dextran FBS (Atlanta Biologicals) one week prior to luciferase experiments to minimize interference from hormones present in media. For luciferase experiments, 100μL of 231GRE cells were seeded at a density of 2.5×10^4^ cells/well in white 96-well plates (Thermo Scientific Nunc, Rochester, NY). The following day, cells were treated, either alone or in combination, with DEX (1×10^−11^-1×10^−5^M), PRED (1×10^−11^-10^−5^M) or MPA (1×10^−11^-5×10^−5^M). Concentrations tested were nontoxic in the MTT assay (data not shown). Untreated (media only), vehicle (DMSO 0.1%) and positive control (100nM DEX) wells were included on every plate. Cells were incubated with chemical treatments for 18 hours at 37°C prior to rinsing with PBS and lysing with 1x cell lysis buffer (Promega, Madison, WI). Luciferase activity was measured using a Berthold Centro XS3 LB 960 microplate luminometer with automatic injection of Luciferase Assay Reagent (Promega). Luminescence measured in DMSO only wells was averaged and subtracted from all values on the plate. Background corrected relative light units were then normalized by dividing by luminescence measured in the 100nM dexamethasone positive control well. Negative numbers were assigned a value of “0.”

### Mathematical models and significance testing

#### Fitting Individual Dose-Response Functions

We recently derived a dose-response function that reflects the pharmacodynamics (PDM) of homodimer nuclear receptors (Webster and Schlezinger 2019). This model assumes a three-step reaction: A+R □ AR □ AR* □ AR**RA. According to this kinetic equation, a ligand (A) reversibly binds its receptor (R) and the ligand-receptor complex (AR) undergoes a conformational change (AR*) that allows homodimers (AR**RA) to form and induce transcription. For a single ligand, the dose-response function is defined by the composite function:

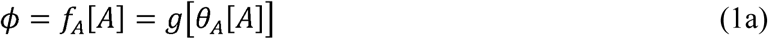

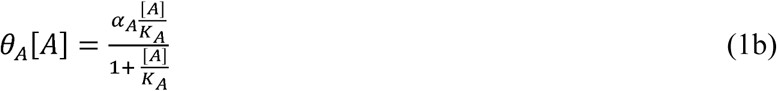

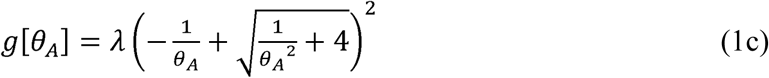

Where *K*_*A*_, *α*_*A*_, and *λ* are all positive parameters. *K*_*A*_ is a macroscopic equilibrium constant and the maximum response for a compound is determined by *α*_*A*_ and *λ*. A ligand independent scaling factor (λ) is included to reflect assay specific variables that influence the measured response (□), such as receptor number. Although these parameters are similar to those obtained by a standard Hill model, they differ in their derivation. It should also be noted that (1c) is slightly different from the equation in Webster and Schlezinger 2019, but is still translatable to the other version via a reparameterization without altering the shape of the dose-response function. Comparisons were made between homodimer functions and Hill functions fit using the drc R package (Ritz et al. 2015).

#### Generalized Concentration Addition (GCA)

One requirement of GCA is specification of an invertible dose-response function for each ligand in the mixture (Howard and Webster 2009). The definition of GCA for two ligands is:

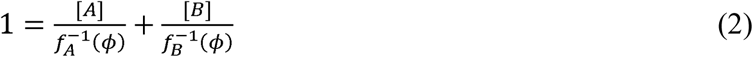

The inverted dose-response functions for ligands A and B are represented by 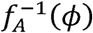 and 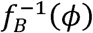. Substituting the inverse homodimer dose-response function:

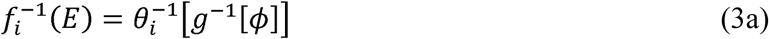

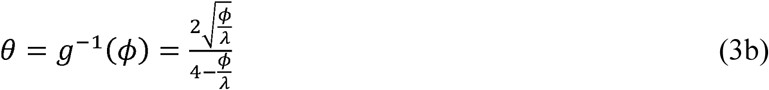

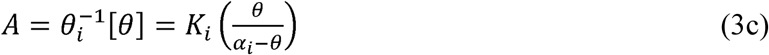

into (4) produces the joint response function of:

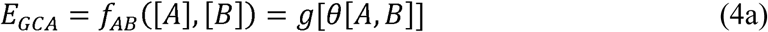

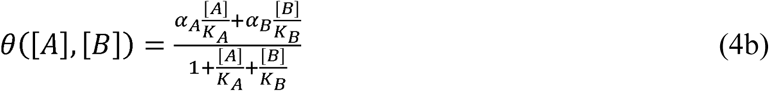

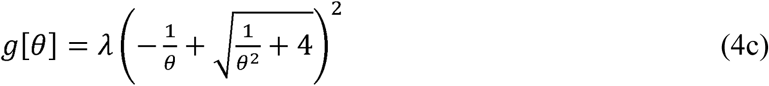

With a common defined for two compounds that differ in *α*_*i*_ and *K*_*i*_.

#### Relative Potency Factor (RPF)

The RPF model assumes that two compounds have dose-response curves with parallel slopes and the same efficacy. RPF is a special case of GCA only when these two assumptions are valid. For RPF, the joint effect of two ligands was predicted using the following equation:

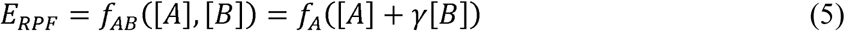

where *γ* is the relative potency of compound B compared to the reference compound A based on their EC_50_ values obtained by fitting a Hill Function for each compound. DEX was used at the reference compound, described by *f*_*A*_([*A*]), since it had the highest potency and efficacy of all tested GR ligands.

#### Effect Summation (ES)

The ES model assumes that the total mixture effect is equivalent to the sum of the individual responses. For ES, the joint effect of two ligands was predicted using the following equation:

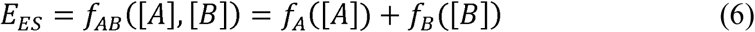

#### Software and Statistics

The R wireframe() function was used to plot the experimental and modeled response surfaces. The nonparametric Wilcoxon rank sum test was used to compare the fit of model predictions to experimental data. This test evaluates whether the experimental and modeled data come from the same distribution. A *p*-value<0.05 indicated a statistically significant difference between the two distributions.

## Results

### Characterizing Independent Dose-Response Functions

The 231GRE cell line was treated with GR ligands, and independent dose-response functions were fit using the homodimer PDM dose-response function (Figure 1). Comparisons were also made between the homodimer PDM dose-response function and Hill functions fit with a Hill coefficient of 1, which assumes a single ligand-binding site (Figure 1). Model parameters for each compound are listed in Table 1. DEX and PRED were both full agonists with similar maximum effect levels. MPA was less efficacious than DEX and PRED, characterizing this ligand as a partial agonist. The Hill and homodimer models had similar RMSE values suggesting that both were comparable. The homodimer PDM dose-response function better characterized the data than the model previously used for receptors with a single ligand-binding site, especially at low concentrations (Figure 1). Therefore, the homodimer PDM function was used to apply GCA to mixtures of GR ligands.

**Table 1.**
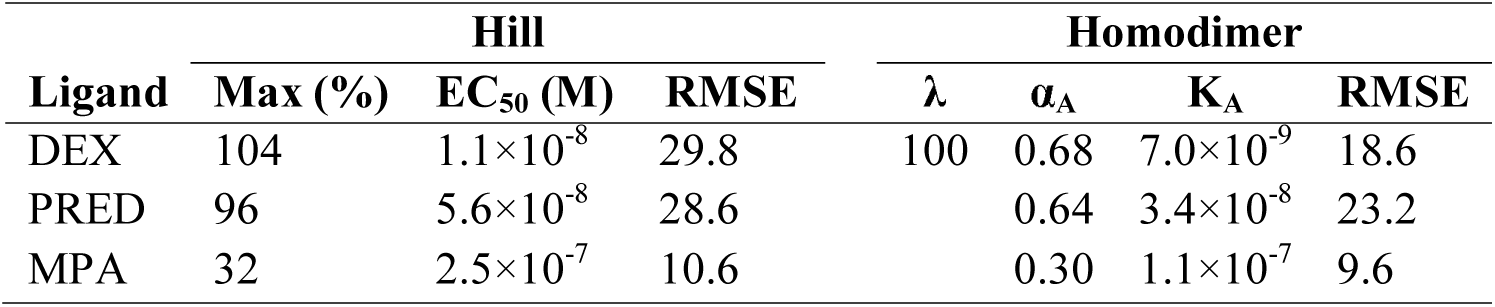
Parameters of the Hill (coef=1) and Homodimer Functions

**Figure 1.**
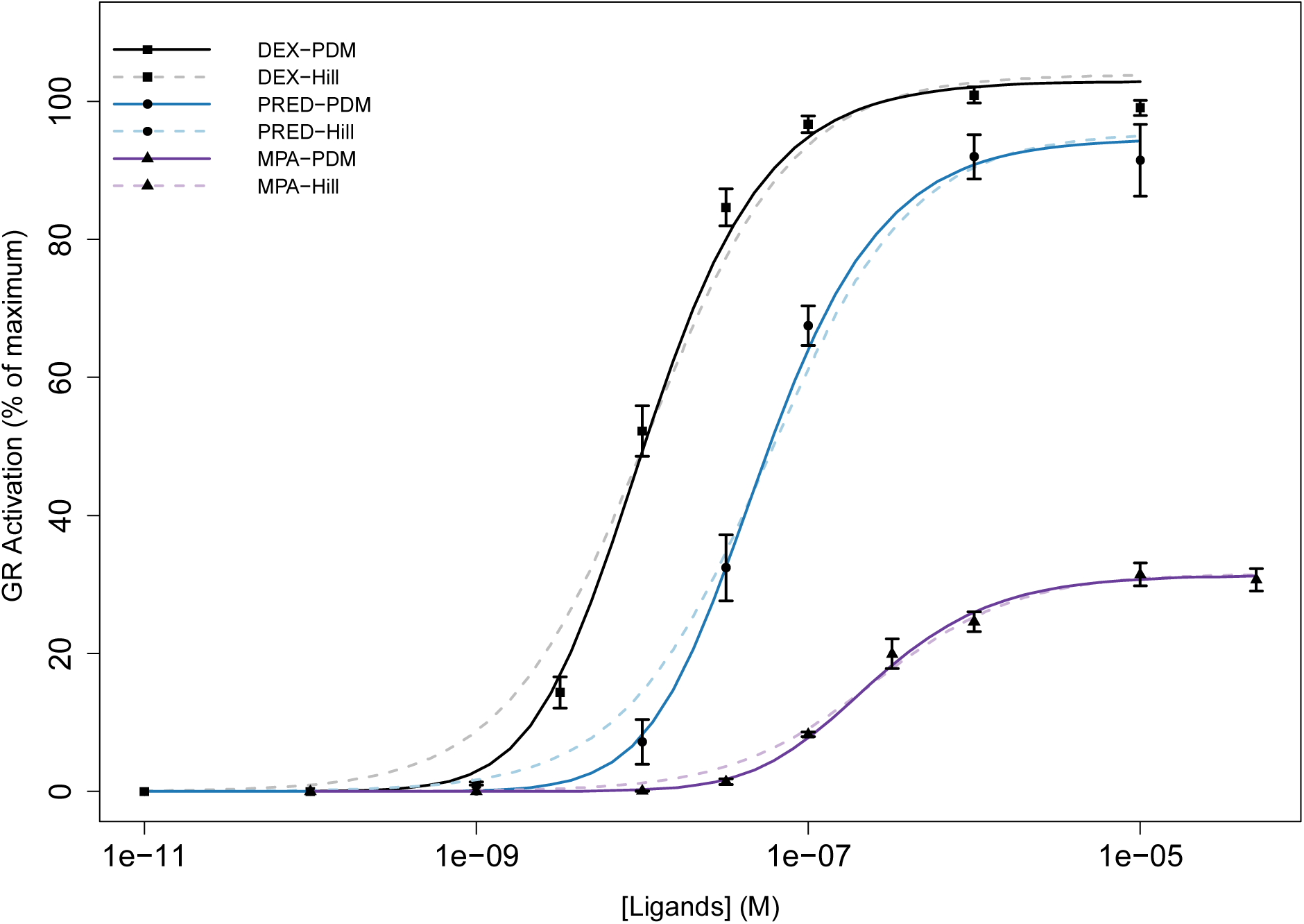
Dose response analysis of GR activation. Reporter data were generated in 231GRE cells treated with Vh (DMSO, 0.1%) or GR ligands for 18 hrs. Dose response data were fit with either a Hill function with a Hill coefficient of 1 (dashed) or a pharmacodynamics (PDM) homodimer function (solid). Error bars represent SEM from three independent experiments (N=3).

### Full Agonist Mixtures

Experimental data for activation of GR by two full agonists were generated using binary mixtures of DEX and PRED. The experimental dose-response surface for two full agonists are show in Figure 2A, with the edge of the box reflecting the marginal dose- responses curves of DEX and PRED. Comparisons were made between the experimental dose-response surface and the joint effects predicted by GCA, RPF, and ES (Figure 2B-D). Non-significant p-values in the Wilcoxon rank-sum test indicated that GCA (p=0.59) and RPF (p=0.35) fit the experimental data reasonably well. However, differences between experimental data and ES predictions approached statistical significance (p=0.08), indicating poor model fit.

**Figure 2.**
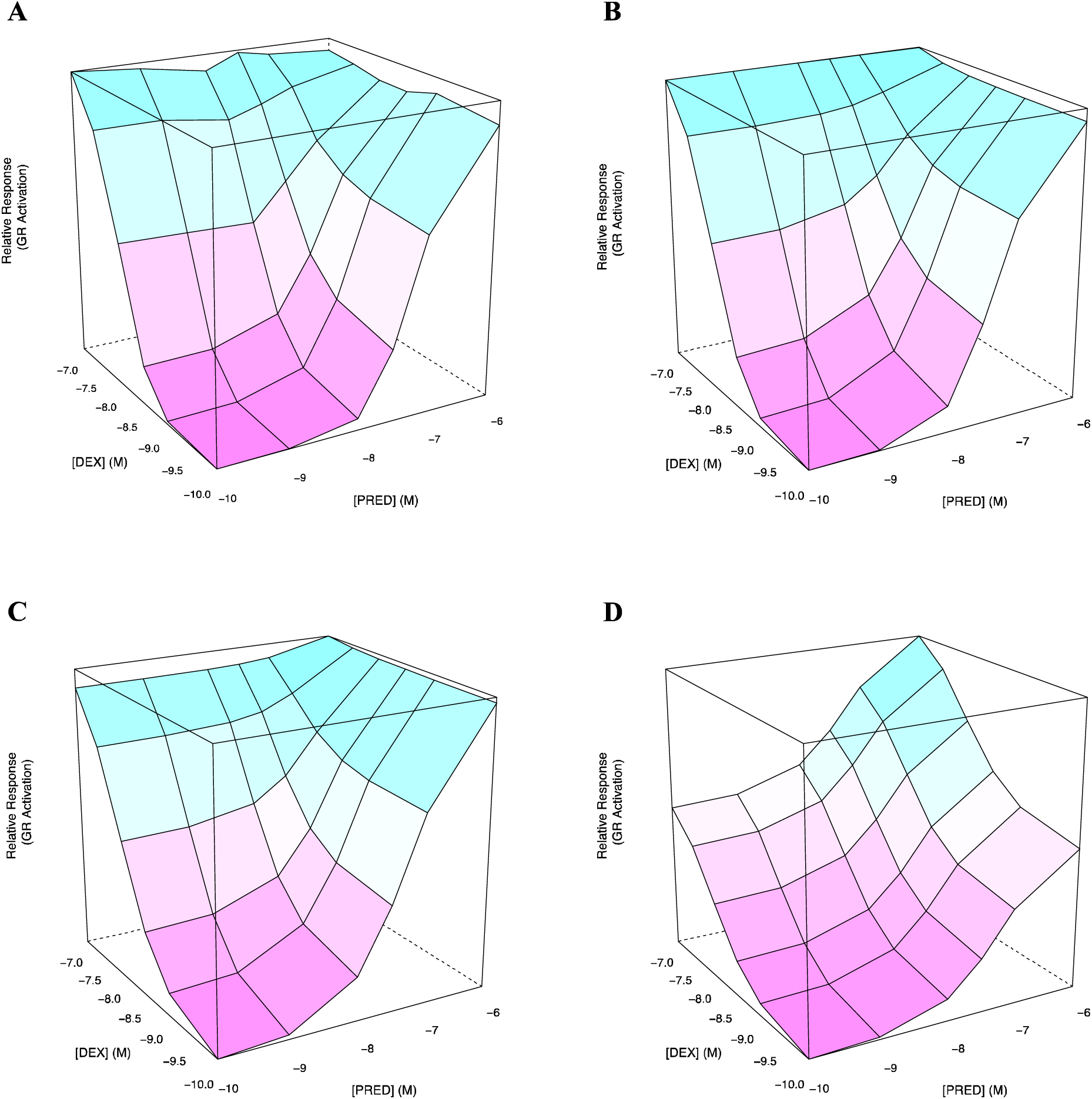
Response surfaces for dexamethasone (DEX) and prednisolone (PRED) mixtures. The experimental data (A) was compared to predictions made by the GCA (B), RPF (C), and ES (D) models. DEX and PRED concentrations are logarithmic. The experimental data surface reflects the mean of three independent experiments.

### Full and Partial Agonist Mixtures

An experimental dose-response surface for a full and partial agonist mixture was generated using binary combinations of DEX and MPA (Figure 3A). MPA increased the GR response at concentrations with lower effect levels. At higher concentrations, where the effect level exceeds the efficacy of the partial agonist, MPA antagonizes the effect of DEX. GCA accounts for this behavior (Figure 3B) and adequately predicted the joint effect of a full and partial agonist (p=0.89). However, predictions made by RPF (p=8×10^−4^) and ES (p=0.05) were a poor fit of the experimental data since they did not adjust for antagonism produced by high concentrations of a partial agonist (Figure 3C, D).

**Figure 3.**
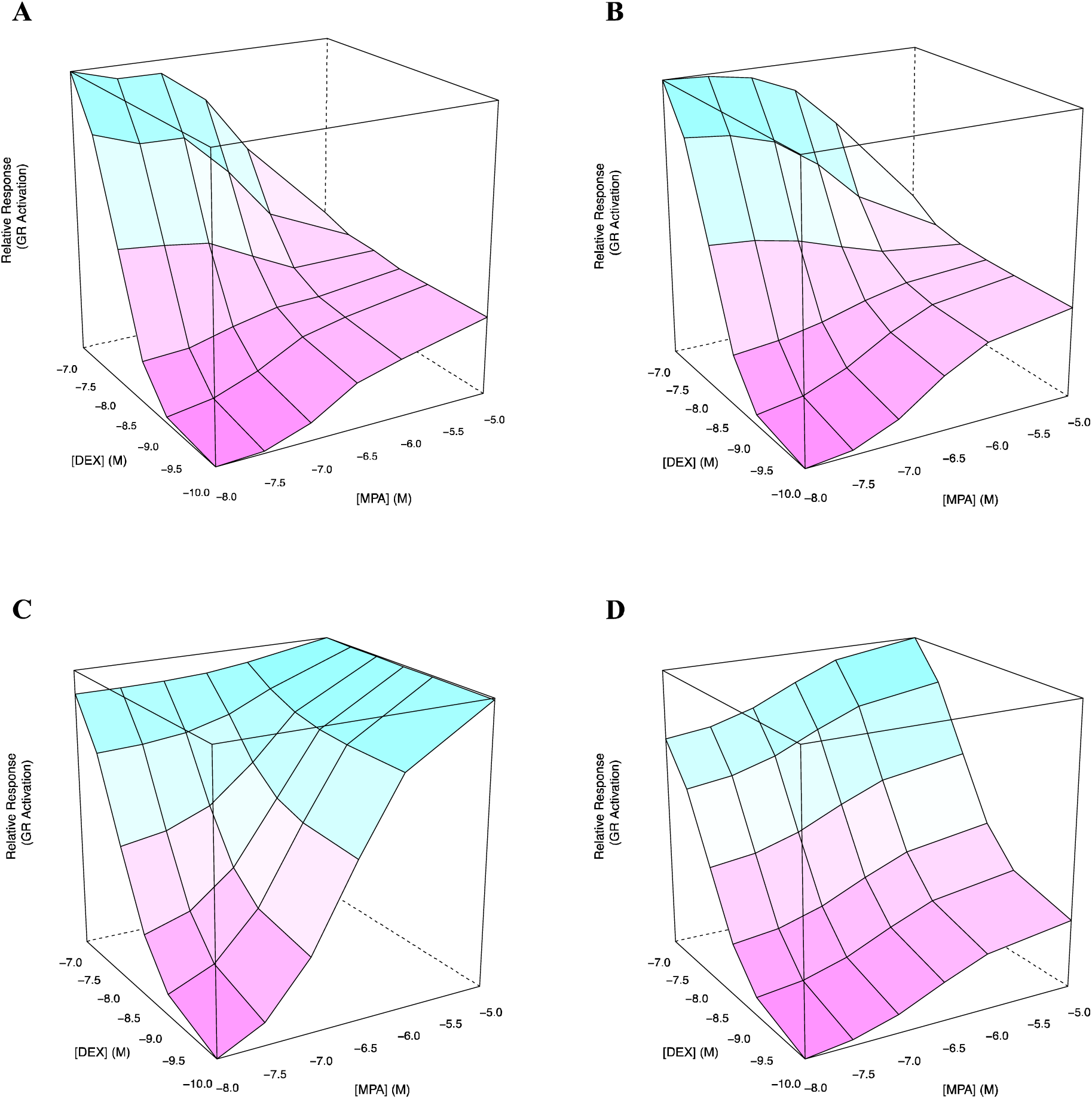
Response surfaces for dexamethasone (DEX) and medroxyprogesterone 17-acetate (MPA) mixtures. The experimental data (A) was compared to predictions made by the GCA (B), RPF (C), and ES (D) models. DEX and MPA concentrations are logarithmic. The experimental data surface reflects the mean of three independent experiments.

## Discussion

This study extends the application of GCA to receptors that homodimerize. We demonstrate that GCA can be applied to ligands that activate GR, a homodimer nuclear receptor. In order to satisfy the requirements of GCA, we used invertible dose-response functions for GR ligands based on pharmacodynamic models for homodimer receptors. We found that individual dose-response data was fit well by the homodimer function. Overall, GCA was the most versatile additivity model. It is able to accommodate mixtures containing either a full or partial agonist. Given that ligands with submaximal efficacy are common for steroid receptors, our extension of GCA to homodimers is an important improvement in the ability to assess and predict the activation of steroid receptors by mixtures of ligands.

The dose-response function used to describe receptors that homodimerize is a fundamental difference between this study and previous work on GCA. For receptors that bind a single ligand, the dose-response relationship can usually be modeled by a Hill function with a Hill coefficient of one (Howard et al. 2010; Watt et al. 2016). This function is invertible, thereby satisfying a critical requirement of GCA. However, an alternative dose-response function is required for ligands of receptors with two binding sites since the Hill coefficient is greater that one, and the inverse Hill function produces imaginary numbers when the response values exceed the estimated maximum value of a compound (Webster and Schlezinger 2019). GR agonists have Hill coefficients greater than 1 since the dose-response function is approximately quadratic at low concentrations. Therefore, we applied GCA to mixtures of GR ligands using pharmacodynamic models for receptors that homodimerize. The composite dose-response function describes binding and activation of the ligand-receptor complex as well as the formation of homodimers from the ligand-receptor monomers. Our model is applicable to multiple biological pathways since the glucocorticoid, mineralocorticoid, androgen, and progesterone receptors are highly homologous and homodimerize (Wahli and Martinez 1991).

Few studies have applied GCA to ligands of homodimer receptors. Brinkmann et al. 2018 demonstrated that GCA more accurately predicted the estrogenic effect of mixtures containing full and partial agonists than CA. The authors applied GCA using our previous approach that assumed a single ligand-binding site for the receptor (Hill function with a Hill coefficient=1). A previous paper also found that GCA, and not CA, predicted the joint effect of binary mixtures containing GR full and partial agonists (Medlock Kakaley et al. 2019). Dose-response curves were fit using four-parameter Hill functions, but they assumed a Hill coefficient of one for use in GCA. While these studies highlight the improvement of GCA over CA in predicting the response of mixtures containing partial agonists, our approach goes one step further by using a more appropriate function to fit dose-response data. The homodimer function met the requirements of GCA and improved prediction accuracy of GR ligands, particularly at low doses. This model also provides information about the underlying biology of an important ligand-receptor interaction.

Synthetic glucocorticoids were used to test whether GCA adequately predicts mixture effects of GR ligands. Nevertheless, this research translates to relevant human exposures. In 2016 the number of prescriptions for prednisolone and dexamethasone in the United States exceeded 4 and 1 million, respectively (Kane, 2018). Furthermore, pharmaceutical glucocorticoids have also been detected in wastewater samples worldwide (Schriks et al. 2010; Kolkman et al. 2013; Macikova et al. 2014; Suzuki et al. 2015; Jia et al. 2016). Humans also endogenously secrete a glucocorticoid called cortisol in response to stress. Hydrocortisone, the synthetic version of cortisol, had 15% lower efficacy for GR than dexamethasone and prednisolone when tested in Tox21(US EPA 2017 Nov 1). Consequently, the response induced by prescribed glucocorticoids could be impaired by high concentrations of circulating cortisol.

There is also evidence that environmental compounds modulate GR activity. Multiple paraben compounds and diethylhexyl phthalate have been shown to behave as partial agonists with low efficacy (Klopčič et al. 2015; Kolšek et al. 2015). However, the majority of tested environmental chemicals antagonize GR activation, some of which include persistent organic pollutants (PCBs, PBDEs and organochlorine pesticides), pyrethroids, metals, and bisphenol compounds (Kojima et al. 2009; Antunes-Fernandes et al. 2011; Zhang et al. 2016; Zhang et al. 2018; Kojima et al. 2019). Therefore, future studies should evaluate whether GCA can predict joint effects of GR antagonists. Additionally, GCA should be applied to more complex mixtures of GR ligands that reflect human exposures.

We used an *in vitro* bioassay to quantify the amount of GR activation induced by ligand mixtures. Our cell line stably expresses a luciferase reporter gene driven by a glucocorticoid responsive element, which produces a response that is directly proportional to the amount of GR activity. This model allows us to characterize the molecular initiating event (MIE), defined as the initial interaction between a chemical and biological target (Ankley et al. 2010). Therefore, evaluating mixture effects of MIEs has broad implications for risk assessment. Future work should examine how predictions made by GCA for MIEs, such as homodimer nuclear receptors, translate to downstream outcomes along the causal pathway.

In conclusion, this study demonstrates that GCA predicts mixture effects of GR ligands. Moreover, our model extends GCA to the broader class of homodimer nuclear receptors (e.g. androgen, mineralocorticoid, and progesterone receptors). We also show that at lower concentrations, the homodimer function describes the dose-response data of GR ligands better than the Hill function previously used for single ligand-binding receptors. Finally, we demonstrate that the GCA model for homodimer receptors adequately fit experimental data of binary GR ligand mixtures, unlike other commonly used additivity models. Future work should evaluate whether GCA can be used to predict mixture effects of pharmaceutical, endogenous, and environmental GR ligands on more downstream biological endpoints. Developing prediction models that reflect these biological processes not only improves accuracy but also informs risk assessment of chemical mixtures.

## Acknowledgements

This project was supported by NIEHS grants P42ES004705 and R01ES027813. The views expressed in this paper are solely those of the authors and do not necessarily reflect those of NIEHS.

## Declaration of Interest

None

## Notes

### Competing Interest Statement

The authors have declared no competing interest.

